# Data quality reporting: good practice for transparent estimates from forest and land cover surveys

**DOI:** 10.1101/399055

**Authors:** Luca Birigazzi, Timothy G. Gregoire, Yelena Finegold, Rocίo D. Cóndor Golec, Marieke Sandker, Emily Donegan, Javier G. P. Gamarra

## Abstract

The need to provide transparent and reliable Greenhouse Gas (GHG) emission estimates is strongly emphasized in the context of international reporting under the United Nations Framework Convention on Climate Change (UN-FCCC) and the Paris Agreement. Yet it is difficult to find specific guidance about what information is really needed to evaluate the quality of the emission factors or activity data used for GHG emission estimates. The most commonly used indicator of the reliability of an estimation procedure (and one of the few indicators explicitly mentioned in the 2006 IPCC guidelines) is the so-called confidence interval, usually at a confidence level of 90% or 95%. This interval, however, is unlikely to be a meaningful indicator of the quality of the estimate, if not associated with additional information about the estimation and survey procedures (such as on the sampling design, measurement protocols or quality control routines, among others). We provide a review of the main sources of error that can have an impact on the precision and accuracy of the estimation of both emission factors and activity data and a list of the essential survey features that should be reported to properly evaluate the quality of a GHG emission estimate. Such list is also applicable to the reporting of national forest inventories and of area estimation of activity data, and includes the case in which confidence intervals are obtained using error propagation techniques.

## 1. Introduction

In order to account for green house gas emissions from the Land Use, Land Use Change and Forestry (LULUCF) sector two approaches are commonly used: *stock change*, when emissions are estimated as a carbon stock difference between two consecutive surveys, and *gain-loss*, whenever emissions are estimated as the product of areas of land use or land use change (aka *activity data*) and the specific carbon coefficient (aka *emission factors*) associated to them (IPCC 2006, Vol.4 Chap.2, GFOI 2016). The Intergovernmental Panel on Climate Change (IPCC) guidelines provide a hierarchical classification of estimation methods, consisting of three levels of methodological complexity, called tiers (IPCC 2003, Chap.3.1.5; IPCC 2006, Vol.1 Chap.1.2). Tier 2 and tier 3 methods, which are considered the most certain and reliable, both rely on sampling up to a certain extent. In order to account for the emissions from LULUCF under these higher tiers, sampling techniques are commonly used for the estimation of both emission factors and activity data. The former are usually derived from in-situ assessments, such as forest inventories or from permanent or temporary experimental plots (Chirici et al., 2011; Köhl et al., 2015), the latter from model-based map classification (wall-to-wall maps) or design-based area estimation through visual, or augmented visual (Bey et al., 2016) interpretation (McRoberts, 2014; FAO, 2016). More advanced methods can involve the use of large-area forest biomass maps, however their use is still limited in the context of GHG inventory and REDD+ reporting (Sandker et al., 2018). In any case, in order for a country to produce reliable higher-tier estimates it is necessary to largely rely on data coming from sampling (IPCC, 2003).

These sample-based estimates are required to be: (1) “accurate, in the sense that they are neither over-nor underestimates as far as can be judged”, and (2) precise, “in the sense that uncertainties are reduced as far as practicable” (IPCC 2003, Chap.5.2; IPCC 2006, Vol.1 Chap.3). Precision and accuracy, as they are defined in the 2003 and 2006 IPCC guidelines, are well known concepts in the literature on probability sampling, where they are usually expressed in terms of variance of an estimator (alternatively called sampling variance) and bias, respectively. As mentioned in the IPCC guidelines, it is worthwhile to mind the difference between the variance of the population and the variance of an estimator. The former provides a measure of how dispersed the values of a population are, while the latter provides a measure of the precision of the estimator used. Even though under simple random sampling there is a direct relationship between these two quantities^1^, they still describe different features, have different applications and should not be confused.

As for any other probabilistic survey, the precision and accuracy of a GHG inventory or of a REDD+ results report can be fully ascertained only in the unrealistic case in which the exact values of interest for all elements in the population are known. In all other cases, precision and accuracy need to be estimated from the sample itself and/or evaluated based on the available information on the studied population, sampling design, statistical assumptions and measurement methods used to obtain the estimates. The so-called confidence interval is one of the most commonly reported indicators of the reliability of an estimation. However, it does not usually include all sources of error in the survey. Confidence intervals typically include the sampling error and in some cases may also partially include some non-sampling errors, such as measurement or model error (cf. Section 2.4 below), but do not measure the bias and other types of non-sampling error (Hanson, 1978). In addition, the confidence interval is a random variable itself and is also estimated from the sample (that is, different random samples will produce different intervals). Several alternative estimation methods may exist to obtain a confidence interval and not all of them are adequate for the specific sampling design adopted (Cochran, 1977; Särndal et al., 1992). In many applications, bias and sampling variance are often treated separately, but they still remain closely interrelated and if the survey is substantially biased, the resulting confidence interval will also be distorted (cf. Raj 1968, Chap. 2.11; Cochran 1977, Chap 1.8). In fact, a point estimate and its associated confidence interval do not reveal whether the reported results are precise and accurate.

Uncertainty is defined in the IPCC (2006) guidelines as the lack of knowledge of the true value of a variable and the word is often used in a broader sense that encompasses both precision and accuracy. The need to estimate and report the uncertainties associated with the estimates is repeatedly stressed in the IPCC guidelines (2003; 2006). They distinguish between uncertainties that are amenable to quantification and others which are non-quantifiable (IPCC, 2006, Vol. 1, Chap. 3). The former, typically including sampling and measurement errors, can be expressed using a confidence interval, while the latter, which may include bias or any type of conceptual or inferential imperfections, cannot. As reported in the IPCC guidelines quantitative uncertainty analysis is performed by estimating the 95 percent confidence interval of the emission and removals. In contrast, non-quantifiable errors, if they cannot be prevented, should be identified, documented and possibly corrected by the compilers. To this end, the guidelines provide eight broad causes of error to be considered by the inventory developers (IPCC, 2006, Vol. 1, Chap. 3) and general guidance on the procedures needed to assess and maintain the quality of the inventory (IPCC 2003, Chap. 4.4; IPCC 2006, Vol.1, Chap.6). However, this recommendation proves to be quite generic and mainly focused on integrity and completeness of the data and does not provide detailed guidance on how to report information stemming from a probability survey.

In the context of a GHG inventory, uncertainty analysis is rather considered as a means to help prioritize national efforts to reduce the uncertainty of inventories in the future, and to guide decisions on methodological choice. They do not set any specific standards concerning which aspects of the survey should be documented. However, it can be beneficial for the reporting Parties to duly demonstrate that their estimates are reliable and that the methods used to obtain them are adequate. From a statistical point of view the confidence interval alone should not be the only quality indicator. Paradoxically, an improvement in survey methods or an increased knowledge of the studied population can lead to wider confidence intervals and give the misleading idea of a decrease in the quality of the estimates. This manuscript aims to provide a comprehensive list of information that: (1) sheds light to reporting parties into confronting this paradox, and (2) should be reported to properly evaluate the quality of a GHGI/REDD+ report estimate for the LULUCF sector. This information is mostly focused on improving the quality declaration of the data, its sources, and the reported estimates as a good practice guidance.

## 2. Guidelines for reporting survey research

Since the 1950s, there have been policies to describe the quality of statistics derived from survey sampling (Statistical Office of the United Nations, 1950; Gonzalez et al., 1975) and many national survey organizations have developed their own quality declaration guidelines (Statistics Canada, 2000; Full et al., 2001; Brackstone, 2003; Office for National Statistics, 2007; Jackson et al., 2013; Brancato et al., 2016). Even though many of those recommendations are certainly useful and applicable in the context of REDD+ or GHG reporting, there are still certain aspects that are somewhat peculiar to the LULUCF sector that are not fully elaborated in those more generic policies. Moreover, the rapid developments in many methodological (and technological) aspects of land use and forest monitoring call for urgent and specific updates in their guidelines. We provide below a description of the main sources of errors which can arise during the estimation of emission factors and activity data from LULUCF sector survey data. For each error source we propose a set of key questions which should be considered by those engaged in reporting or reviewing survey results. The answers to such questions will constitute the essential body of information that should be reported to allow reviewers, reporting Parties and practitioners to properly evaluate the quality of a GHGI/REDD+ report estimate. When the emission factors and activity data have been estimated through independent surveys the answers should be provided for each of them. Guidance for the reporting of the combined uncertainty of emission factors and activity data using error propagation techniques is provided in section 2.3.

### 2.1. General information about the survey

This section aims to provide general information about the survey, including a description of the population sampled, the data collected, the methods of measurement and the sampling design adopted. We assume that readers already have some knowledge about the basics of sampling.

#### 2.1.1. Information about the sampled population

The term *sampled population* denotes the “aggregate from which the sample is chosen” (Cochran, 1977). The population is composed of *elements*, to which one or more variable of study are associated (Särndal et al., 1992, Chap. 1.2). The sampled population is identified at the planning phase of a survey and should be defined in such a way that there cannot be any ambiguity about whether or not an element is part of the population (Köhl et al., 2006). When sampling for emission factor or activity data for the LULUCF sector the sampled population is often defined as a geographic area. In this case, the population includes all locations that have non-zero probability of being included in the sample. In National Forest Inventories (NFIs) the population typically corresponds to the whole country area or, in some cases, to the area of the country that is considered forest. When subnational surveys are carried out, the sampled population may correspond to a specific administrative unit or to a particular ecological zone.

When ground-surveys are carried out it is possible to define the population as a continuous areal frame. That is, it comprises an infinite number of spatial locations (Gregoire and Valentine, 2008; Köhl et al., 2006). In remote sensing applications, in contrast, the population is often defined as a finite set of non-overlapping spatial units that form a partition of the region of interest, typically pixels, block of pixels or polygons. In this case, the choice of the type of spatial units that tesselate the population has an impact on the survey estimates (Stehman and Wickham, 2011) and should be adequately described.

**Set 1 of key questions: Sampled population**

a. What is the population from which the sample is chosen?
b. If the sampled population is defined as a geographic area, are you able to provide a map of it?
c. Is the sampled population defined as a finite set of discrete spatial units? If so, which ones? are they uniform? what is their area?

#### 2.1.2. Information about the target population

The term *target population* denotes the population about which the information is wanted. Similarly to above, when sampling for emission factors or activity data for the LULUCF sector the targeted population is often defined as a geographic area (McRoberts et al., 2015). This may or may not coincide with the sampled population. In fact, in GHG inventories the population of interest is often a sub-group of the sampled population, created after (and independently of) the sample selection, such as a specific land use, forest type or climatic zone (cf. Section 2.2.3 below). Figure 2 provides a visual representation of some cases in which target and sampled population differ.

**Set 2 of key questions: Target population**

a. What is the population for which we want to estimate emissions/removals?
b. For which time period do we need the emissions/removals?
c. If the target population is defined as a geographic area, are you able to provide a map of it?
d. If the target population is not defined as a geographic area, are you able to provide a list of the elements that compose it?

#### 2.1.3. Sampling selection

There are many existing approaches to select a sample from the population. The choice of the sampling method has an important effect on the quality of the estimates and should be carefully described. In NFIs or in land cover area estimation the sample is usually composed of a set of locations selected from a continuous areal frame (such as a geographical region). The unit of area that is observed is often called a plot. A sampling unit can also be composed of a group (cluster) of subplots located near each other (Kangas and Maltamo, 2006) and/or of one or more nested smaller subplots (Kohl et al., 2006).

**Set 3 of key questions: Sampling selection**

a. Is the survey based on a probability sample^2^?
b. What sampling design has been used?
c. What was the planned size of the sample?
d. What is the size and shape of the sampling units?
e. Is the sampling unit composed of a cluster of subplots?
f. Is the sampling unit composed of one or more nested smaller subplots?
g. Was the sample selected following stratified sampling?
h. If so, which are the strata? how were they constructed? What is their size?
i. If so, when the strata are defined as geographical areas, are you able to provide a map of them?
j. Repetition: Is the survey isolated or is it part of a series of repeated surveys? If so, what is the proportion of samples that were repeated?

#### 2.1.4. Data collection, labeling, coding and editing

Once the sampling design is established, the data are collected according to the prescribed measurement protocol, coded and entered into a database. In NFIs, the protocol for collecting the data in the sampling units is usually described in a field manual, and in remote sensing applications (such as accuracy assessment of maps) the procedure used to collect information from each sampling unit is often referred to as *evaluation protocol* (Stehman and Czaplewski, 1998). The choice of the data labeling and of the data management system has a large impact on the precision and accuracy of the results. While conceiving the survey, a particular attention should be put in the definition of categorical variables, such as for example land cover. In order to assign each element to a certain land cover class it is necessary to have a consistent and complete land cover classification system. The classification system should be defined in such a way that each land cover element can clearly be assigned into one and only one land cover class. In remote sensing applications, the set of procedure to assign a classification to each sampling unit is often called *labelling protocol* (Stehman and Czaplewski, 1998) and should be adequately described.

**Set 4 of key questions: Data collection and processing**

a. Which attributes have been observed in the sampling units?
b. What was the measurement protocol used to measure the variables of interest?
c. Has a written field manual or evaluation protocol been produced? Can the party provide it?
d. Can the party provide a clear and unambiguous definition for each class of categorical variables in the survey, including a land cover classification system (if any)?
e. Has the classification system (if any) been modified during the implementation of the survey? If so, the final estimates? Did the party put in place any system to account for this change?
f. Has the field measurement protocol been changed during the implementation of the survey? If so, how does that impact the final estimates? Did the party put in place any system to account for this change?
g. How were the data stored and processed?

### 2.2. Information about error sources

This section aims to provide specific information about the potential sources of error in the survey which require description and analysis. In the literature on survey sampling the sources of error are typically divided into the two broad categories of sampling and non-sampling errors. Abiding by the terminology used in Särndal et al. (1992), non-sampling errors can be further divided into 1) *errors due to nonobservation*, when it is not possible to obtain data from parts of the population of interest and 2) *errors in observations*, when the recorded value of the sampled element differs from its real value. The former includes frame imperfections and non-response issues, the latter measurement and processing errors. All these sources of errors can affect both the precision and the accuracy of the estimates and will be discussed in detail in the next sections, all enumerated and linked to corresponding sets of key questions (Fig. 1). A complete theory of non sampling error has not been elaborated yet (cf. Särndal et al., 1992, Chap. 14.6) and the schema proposed in Fig. 1 does not intend to provide a exhaustive and consistent classification. Moreover, Figure 1 does not define in detail all sources of uncertainty that can arise during the estimation of emission factors or activity data. The aim here is rather to provide a conceptual framework to group the errors into major broad categories. The workflow for satellite data processing or, for example, for tree field measurement, involves multiple steps that are not explicitly mentioned in Fig. 1 (such as satellite sensor calibration or tree biomass allometric model selection). All these steps, however, can always be classified into one of the broad categories mentioned above. More detailed lists of the error sources typically arising in the processing chain to calculate forest emission factors/activity data are provided elsewhere (Hill et al., 2013; Sandker et al., 2018).

**Figure 1:**
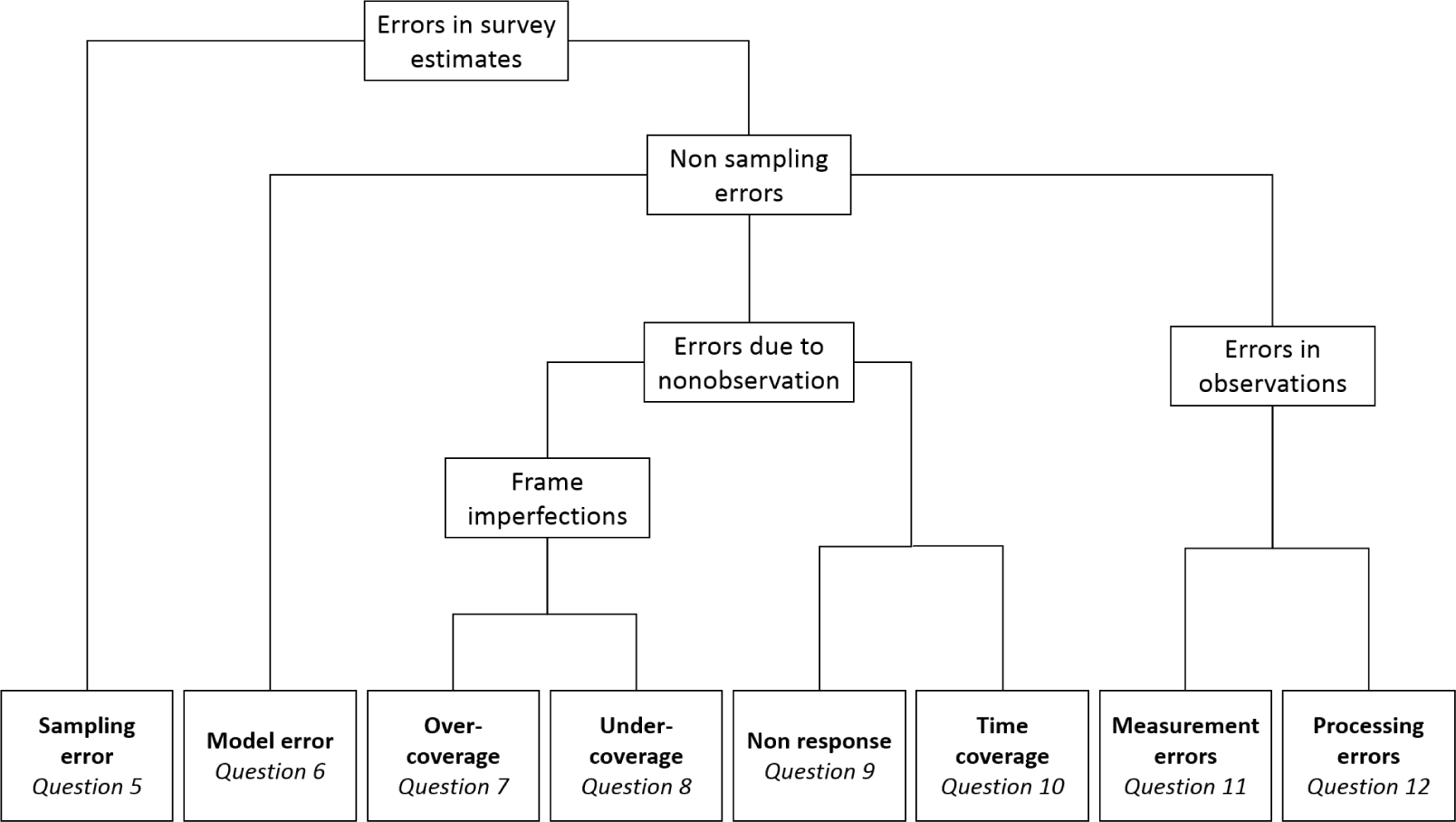
Categories of potential error sources in the LULUCF sector. Broad categories of error sources in surveys sampling and a reference to the corresponding set of key questions in this paper. This schema is aimed to provide a practical framework for reporting survey results.

**Figure 2:**
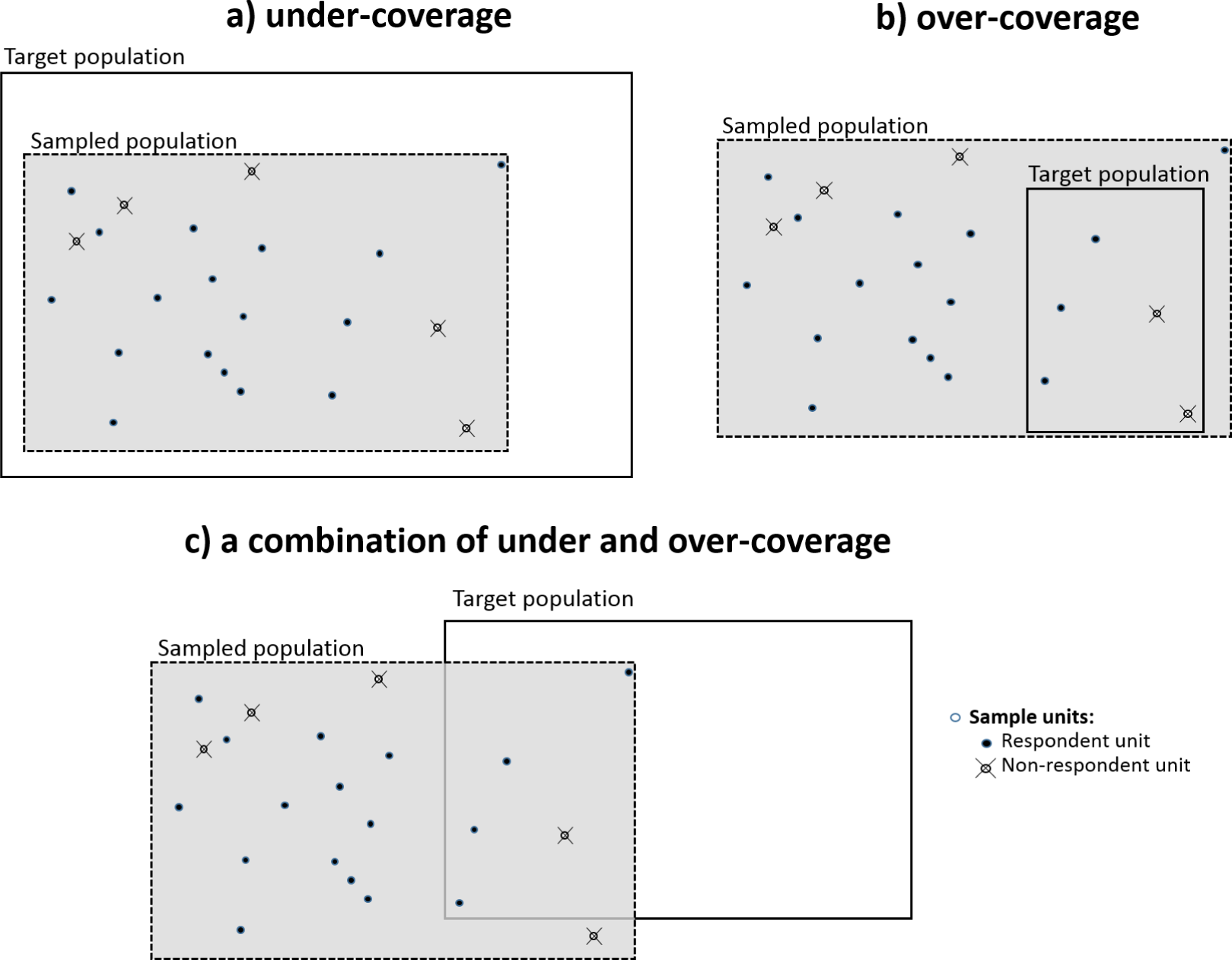
Non-response and frame imperfections. Three cases in which the target population (delimited by the solid line) does not coincide with the sampled population (the gray area delimited by the dotted line): a) the sampled population is smaller than the target population and entirely included in the target population; b) the target population is smaller than the sampled population and entirely included in the sampled population; the target population is not entirely included in the sampled population and vice versa. Circles represent the sample units and include both the non respondent units (crossed circles) and the respondent ones (black circles).

There are two essential complementary approaches to deal with each of the error sources: 1) measures put in place to prevent (or possibly avoid) the error before it occurs (Duvemo and Lämås, 2006; Gasparini et al., 2009) and 2) apply methods to properly account for the error once it has occurred (Pollard et al., 2006; Ferretti et al., 2009; Gormanson et al., 2018). A complete assessment of any survey result cannot be done without a thorough analysis of these two aspects. Hence, as a matter of transparency, reporting parties should take care to describe both of them. In the following sections for each error we provide recommendations to ensure that both aspects are duly included in the reporting.

#### 2.2.1. Sampling error

The sampling error denotes the error caused by the fact that only a subset (a sample) of the population is measured. Even if no error is made in measuring or processing the data, it is still evident that the estimates based on the sample will differ from the real population values. On the other hand, it is intuitive that different samples of the same population will provide different estimates (unless the population is composed of identical elements). The *variance* of an estimator (or sampling variance) provides a measure of the sample-to-sample variation and the *bias* of an estimator is the difference between the real population value and the average of all possible sample estimates. Under many sampling designs it is possible to provide a quantitative unbiased estimate of the sampling variance. For any given estimator of a population parameter many statistics manuals also provide: 1) the formula for the *estimator variance* (in practice an unknown quantity which depends on the complete set of population values) and 2) a formula to estimate unbiasedly the estimator variance from the sample data (cf. Särndal et al., 1992, Remark 2.8.2).The latter is the one ordinaryly used to compute the sampling error. Hence, in most of the cases the sampling variance falls within the IPCC category of errors amenable to quantification and can be expressed using a confidence interval. In practice, confidence intervals calculated on the sampling variance are the most commonly reported indicators of the reliability of an estimation. Larger sampling variances will result in wider confidence intervals, hence in an overall decrease of precision. The bias of the estimator, conversely, is often not quantifiable in practice and its magnitude can only be inferred from the sampling design, the estimators used and the population parameters being estimated.

Sampling error will always be present, unless the population is enumerated in its entirety. Typical examples of surveys sampling used in REDD+ and GHG reporting are national forest inventories (for forest emission factors) or area estimation (for land use/land use change activity data) (Olofsson et al., 2014). For a given sampling design there may exist several alternative estimators, each one having different statistical properties. The reporting Parties should take care in selecting the best estimator, where best here means having small variance and being unbiased (or approximately unbiased). The properties of the estimators under the most common sampling designs have been thoroughly investigated, so that it is usually possible to estimate their variances and to ascertain whether or not they are unbiased (or approximately unbiased). A plethora of manuals has been dedicated to the theory and practice of sampling methods, including the renowned texts of Cochran (1977)) and Särndal et al. (1992). de Vries (1986), Kangas and Maltamo (2006), Köhl et al. (2006), Gregoire and Valentine (2008) and Mandallaz (2008), among others, provide a more detailed review of sampling strategies for natural resources and forest inventories.

**Set 5 of key questions: Sampling error**

a. Which estimator has been used to estimate the emission/removal? Can the party provide the mathematical formula used?
b. Has the variance of the estimator been estimated? If so, which estimator was used to compute it? Can the party provide the mathematical formula used?
c. Is the estimator for the population parameter and its variance estimator unbiased (or approximately unbiased) under the sample design adopted in the survey?

#### 2.2.2. Model error

Many of the variables of interest in REDD+ or GHG reporting are not directly measured in the field but estimated from other observed variables. This is the case of tree biomass, carbon or volume, usually predicted using allometric models with one or more easy-to-measure explanatory variables (such as tree height or diameter at breast height). The fact that the variables of interest are predicted and not measured adds additional uncertainty to the estimation process and is likely to result in a decrease of both precision and accuracy. Model error is used here to denote the error between the real element value (such as the aboveground biomass of a certain tree) and the value predicted by the model (assuming no measurement or precessing error are made). Two main sources of uncertainty contribute to this type of error: the uncertainty in the estimation of model parameters and the random model residuals. In large area surveys such as NFIs, however, the latter is typically very small (Chambers and Clark, 2012) and only the error in the estimation of the model parameters contributes significantly to the total error (Ståhl et al., 2016).

Ideally, model development constitutes a phase of the survey sampling itself and data needed for model predictions are sampled using the NFI design. In this case (and if an adequate sampling design is used) it can be possible to demonstrate that the model prediction is unbiased (or approximately unbiased). In practice, the application of models developed before and independently of the survey sampling is common in NFI and GHG reporting. Regional models, or models constructed by global macro ecological zone, such as the pantropical biomass regression of Chave et al. (2014), are extensively used worldwide. The implicit (and often critical) assumption here is that the population for which the model was developed is very similar (if not identical) to the population of which we want to report the emissions/removals (Cunia, 1986). If this assumption does not hold, the model predictions are very likely to be biased and such a bias will propagate throughout the whole estimation process. Some authors also quantify the error in model choice, that is, the uncertainty due to the fact that more than one models exists in the literature and the reporting party does not know how to arbitrate between them (Chave et al., 2004; Picard et al., 2015; Duncanson et al., 2017). Reporting parties should pay attention to justifying the choice of the models used and possibly demonstrate their applicability to the target population. In theory, the model bias can be quantified using specific validation techniques based on national data (Claeskens et al. 2008, pp. 172 and 232; Woodall et al. 2010). In addition, previously constructed models available in the literature often do not provide the key statistics needed to compute the model error. In some cases, methods exist to estimate the model error in absence of the covariance matrix (Magnussen and Carillo Negrete, 2015) or through simulation of pseudo-data (Wayson et al., 2015). Methods for accounting for model errors in NFI when model data are sampled using the NFI design are presented in Cunia (1986) and Ståhl et al. (2014). Monte Carlo simulations are also often used to account for the model error (see also section 2.3 below) and specific statistical software or packages have been developed for this purpose. Réjou-Méchain et al. (2017) provide a Monte Carlo algorithm in R (R Core Team, 2016) to account for the error in the parameters of the model of Chave et al. (2014). Specific guidelines for documenting and reporting tree allometric equations are provided in Cifuentes Jara et al. (2015).

Model errors also abound in satellite-derived data. Image preprocessing is necessary to account for sensor, solar, atmospheric, and topographic effects; however, it can increase the potential to introduce error (Kennedy et al., 2009). Supervised and unsupervised image classification errors are particularly pervasive under the LULUCF sector approaches for the generation of activity data (Potapov et al., 2014) and tend to introduce considerable bias (Hill et al., 2013; Olofsson et al., 2013, 2014), filtering choices, spatiotemporal averaging, interpolation and extrapolation, among others, can contribute to increased uncertainties from satellite-derived data when linked to sampling through the use of training datasets (Hill et al., 2013).

**Set 6 of key questions: Model error**

a. Was the variable of interest observed or estimated using a model?
b. If so, what model was used?
c. What auxiliary variables have been used to estimate the variable of interest? How have they been collected?
d. Were the auxiliary variables available for all population elements?
e. Was the model developed independently of the survey (i.e. selected from models already published in the literature)?
f. How was the model selected and how can the party ensure that the population for which the model was developed is similar to the target population?
g. If the data used to develop the model were collected within the survey, can the party provide a description of the survey design and of the model fitting method?
h. Was the error in model parameters estimated? how?

#### 2.2.3. Frame imperfections

Ideally, the sampled population should coincide with the population about which emission factors or activity data are wanted. Any differences between these two populations may constitute a departure from the ideal conditions for the probability sampling approach and should be accounted for (Lesser and Kalsbeek, 1999). Figure 2 shows three examples of frame imperfections that can arise during the estimation of emissions/removals from LULUCF sector survey data.

#### 2.2.4. Frame imperfections: under-coverage

If the population that has been sampled is only a subset of the population of which we want to estimate emissions/removals, the properties of the estimates will be affected. In the literature on survey sampling this issue is often referred to as under-coverage and it is very likely to result in some bias in the estimates (Särndal et al., 1992; Särndal and Lundsträm, 2005). In the context of the REDD+/LULUCF sector this can occur if the Party wishes to report emissions/removals at national level but without the use of data from a national level survey; instead sampling only limited areas of the country such as a specific region, ecological zone, forest type or ad hoc research plots. Special estimation techniques (such as weighting or imputation) can be used to adjust for under-coverage but they often require the use of auxiliary variables and/or strong assumptions regarding the population of interest. Even if no advanced modeling techniques are used, a party should still duly report the assumptions and the expert judgments on which they have relied to correct for under-coverage.

**Set 7 of key questions: under-coverage**

a. Is the area over which we want to report the emission/removals entirely included in the area that has been sampled?
b. If not, how did the party ensure the estimates are representative of whole target population?

#### 2.2.5. Frame imperfections: over-coverage (domain estimation)

Specific statistical techniques should be used in case the population for which we want to report the estimate is smaller than the population sampled, that is, the opposite of the case described in section 2.2.4. This can occur frequently in REDD+/GHG reporting, whenever data from national or subnational surveys are used to obtain different emission factors/activity data for a set of subpopulations of interest (such as by forest type, ecological zone, district, etc.). In the literature on survey sampling these subpopulations of interest are also called domains. The problem stems from the fact that the number of samples falling into a certain domain is random (i.e. it is not controlled by the inventory designers) and, most likely, small. This can result in a decrease in precision (that is, wider confidence intervals) and, if the right statistical approach is not used, in a bias in the estimate. A detailed review of basic estimation methods for domains is provided in Section 10.3 of Särndal et al. (1992). A more specific discussion about domain estimation in the context of GHG and REDD+ reporting is presented in Birigazzi et al. (2018). If the sample size of a certain domain is particularly small (which is likely to occur when also the domain size is small, such as a very small administrative unit or a very rare ecosystem), the estimation may require the use of ancillary data or model-based inference, which may in turn compound uncertainties with the model errors previously discussed. In the literature this issue is often referred to as small area estimation (Schreuder et al., 1993; Rao and Molina, 2015).

**Set 8 of key questions: over-coverage (domain estimation)**

a. Is the area over which we want to report the emission/removals (the target population) smaller than the area that has been sampled and included in it?
b. If so, has the party used any domain estimation techniques? Which ones?

#### 2.2.6. Non-response

Non-response is the term used in statistical literature to refer to the failure to measure some of the units in the selected sample (Cochran, 1977). In national forest inventories this typically occurs whenever, for example, some sample plots cannot be accessed by the field crews, and the tree variables cannot be measured. In the context of remotely sensed area estimation it can occur if some images are not available or masked by clouds and cannot be interpreted. Non-response in remote sensing applications can also be due to a malfunction in the satellite data collection mechanisms, such as the failure of the Scan Line Corrector of Landsat-7 (Markham et al., 2004), which results in gaps in the imagery. Both the variance and the bias are likely to increase together with the non-response rate. The fact that actual sample size turns out to be smaller than what was originally planned can result in an increase in variance (i.e. a wider confidence interval). On the other hand, the bias can derive from the fact that non-responding elements may be systematically different from the responding ones (Särndal and Lundstr öm, 2005). In general, the wider the difference in terms of average values between the non-respondents and the respondents, the bigger the bias. Methods exist for dealing with non-response both before and after the data collection. The latter, which are often referred to as non-response adjustment may include the use of auxiliary data, as in the case of weighting and imputation methods, or include an additional subsampling of the non-respondents. General principles to assist the estimation in the case of non-response are provided by Särndal and Lundström (2005).

**Set 9 of key questions: Non-response**

a. Did the party put in place any measure for the prevention or avoidance of non-response before the data collection? If yes, which ones?
b. How many of the selected sampling units have proven to be not measurable/not accessible?
c. Which are the main causes for the non-response? Are there any reason to believe that non-responding elements may be systematically different from the responding ones?
d. Did the party adjust the estimate to overcome the fact that not all the sampling units have been measured/accessed? If so, how?

#### 2.2.7. Time coverage issues

A survey is typically carried out in different consecutive phases. Sample selection, data collection, data processing and the estimation obviously do not occur at the same point in time. Since the attributes of interests of the population elements are likely to change over time, it is fundamental to specify the time point in which such attributes were observed. On the other hand, the reporting parties need to report emissions/removals for a specific time period. In the literature on survey sampling this is often called *reference time point for the target population*. The lag between the moment in which the variables are observed and the reference time point for the target population should be as short as possible to reduce the potential time coverage error (Särndal and Lundstr öm, 2005). In case this time lag is particularly large it is possible to use specific interpolation and extrapolation techniques and develop time series for the variables of interest (cf. IPCC, 2006, Vol. 1, Chap. 5).

**Set 10 of key questions: Time coverage issues**

a. When was the data collection carried out? What is the time period in which the variables of interest were observed?
b. Is the period over which we want to report emissions/removals included in the data collection period?
c. If not, how did the party ensure that estimates are representative of the reporting time period?

#### 2.2.8. Measurement error

The measurement error denotes the difference between the real element value and the values that are measured during data collection. In the context of an NFI this typically includes the errors in measuring tree dendrometric parameters (such as tree height, diameter and species, among others), and in the remotely sensed estimation of activity data, such as through visual interpretation of aerial or satellite imagery, it may encompass the interpreter error. The spatial uncertainty associated with the location of the observations (aka *position error*) is another example of measurement error which can have considerable impact on the estimation, especially in the remote sensed assisted estimation of land use change (Cressie and Kornak, 2003). Measurement errors affect both the bias and precision of the estimators. Measurement error from human interpretation can be reduced by ensuring adequate training of the field operators or interpreters and by making use of more accurate measuring instruments. There exist methods to account for the measurement error after the data collection. They rely on the use of repeated measurements, re-measurements using more accurate devices or on the development of more complex measurement models. Cochran (1977, Sect. 13) and Särndal et al. (1992, Sect. 16) discuss general methods for dealing with measurement errors. Measurement error computation in tree height (Larjavaara and Muller-Landau, 2013), or among-interpreter error through augmented visual interpretation (Bastin et al., 2017) are common examples. A recent paper by McRoberts et al. (2018) investigates the effect of the interpreter error on remote sensing-assisted estimators of land cover class proportions. Measurement errors purely rooted in satellite products may also include sensor calibration and degradation, irradiance variation, radiometric resolution, signal digitization, sensor drift or athmospheric attenuation and path radiance, and will further introduce systematic errors in later processing and modelling phases (Curran and Hay, 1986; Hill et al., 2013).

**Set 11 of key questions: Measurement error**

a. Which measuring instruments were used to measure the variables of interest? Is there any information available about the nominal precision of the measuring instruments used?
b. Which instruments were used to record the geographical location of the observations? Is there any information available about the nominal precision of the instruments used?
c. Can the party provide the precision with which each measurement is taken (for example, diameter at breast height to be recorded in cm to the nearest 0.1 cm)?
d. Did the party put in place any measure for the reduction of the measurement error (including position error)? If yes, which ones?
e. How many observers/field teams have been employed in the survey?
f. which satellite products were used to estimate activity data? what are the product specifications?

#### 2.2.9. Processing errors

These include the errors occurring during the coding, editing and processing of the data. In NFIs this can encompass the mistakes made while entering the data in the field forms and/or in the database. Since thousands of elements (such as trees, or land cover sample units) are usually observed in large area environmental surveys, it is almost inevitable that some incorrect values are recorded. The number of potential processing errors is such that it would be difficult to provide a complete list. This will depend on the survey type, on the data processing chain and on the nature of the different variables collected. In NFIs, in the context of GHG or REDD+ reporting, dendrometric variables (such as tree height, diameter, species, among others) are of great importance and the reporting parties should ensure all of them have been carefully assessed for quality. Processing errors affect both the precision and accuracy of the results. Given the multifarious nature of this source of error, it is difficult to provide a measure of its impact on the estimates, but it can certainly prove to be extremely relevant when no preventive or corrective measures are taken. The use of electronic tablets instead of paper forms (possibly associated with validation rules to warn the users whenever potentially erroneous values are entered) is an example of a measure to prevent data entry error in the field. Protocols for data cleaning in the office may include routines for the identification of outliers or missing data using graphical or statistical approaches. Methods for filling missing data and correcting incorrect values may include modelling or interpolation techniques and should be carefully described by the reporting parties. An overview of good practice for data entry and data quality control for NFI is provided by Morales-Hidalgo et al. (2017).

**Set 12 of key questions: Processing errors**

a. Did the party put in place any measure(s) for the prevention of data entry error in the field and in the office? If yes, which?
b. Which methods were used for the identification of invalid or aberrant values after the data were entered?
c. Which methods were used for correcting missing data and outliers?

### 2.3. Total (propagating) errors

International reporting of errors to provide a final uncertainty estimate in the context of REDD+ and GHGI may involve the propagation of some or all of these errors as the total error, in some cases combining both emission factors and activity data uncertainties. Reporting parties are so far free to choose which errors to report, although typically they report the sampling error as a minimum. It is trivial, but very important to recall that the more errors accounted for, the wider the total error reported. In the simplest scenario, often linked to Tier 1 approaches, reporting parties will delve into the standard formulas for the propagation of error (aka delta method or Taylor series) (Ku 1966; IPCC 2006, Vol. 1, Chap. 3). These formulas, much older than some more computational approaches, such as re-sampling, rely often heavily on assumptions of distributional symmetry, trivial covariance structures, or model misspecifications. However, they are in most cases standard formulas easy to implement. Monte Carlo approaches are typically used under higher Tiers estimation methods (Birdsey et al., 2013). They rely on repetitive random draws of values based on probability density functions for emission factors and/or activity data. By their very nature, they can in many cases propagate errors without assuming particular distributions in residuals and can tackle better correlations between variables or situations where widespread distributions are the norm (IPCC, 2006). However, they must be carefully developed and entail larger computational requirements. In some cases, they may inherit some simple assumptions from purely design-based estimators or traditional error propagation approaches. As a result, uncertainty outputs from Monte Carlo approaches are typically provided in the form of probability density functions, or more simply, as a result of the likely asymmetric distributions resulting from the computation, they can be reported through the resulting quantiles (McMurray et al., 2017). Careful use of assumptions regarding correlations between variables and the use of sensitivity analyses cannot be forgone (Heath and Smith, 2000). Monte Carlo outputs can often be re-sampled to obtain the final confidence intervals. Rubinstein and Kroese (2008) provide a good textbook to delve into the topic. Although not too often used so far, example calculations in the context of REDD+ are gaining momentum (Pelletier et al., 2011; Köhl et al., 2015; McRoberts and Westfall, 2016).

Currently not included as a calculation option by IPCC, Bayesian approaches use a form of Monte Carlo computation, but rely on prior information to obtain uncertainties (Fox et al., 2011; Molto et al., 2013). Un-certainty comes in the form of the so-called credible intervals. It is unclear, however, how these uncertainties should be reported within the context of this manuscript.

**Set 13 of key questions: error propagation**

a. Did the Party propagate errors to establish the total error?
b. Were all the errors symmetric or normally distributed? If not, how were those errors reported?
c. Which assumptions were taken in regard to the existence of correlations among the different errors?
d. If errors were propagated, which method was used and how was it justified?
e. If the total error followed an asymmetric distribution, how was it reported?

### 2.4. Good practice for reporting confidence intervals

Confidence intervals (CI) are meant to provide a measure of how well the parameter of the population has been estimated and are one of the few indicators of the quality of the estimate explicitly mentioned in the IPCC guidelines. Even though they usually include only the sampling error, they may also include one or more of the other errors indicated in Figure 1 (cf. Fig. 3). It is evident that as more sources of error are included, the wider the confidence interval will be. In addition to that, the confidence intervals are also a function of the confidence level: the larger the confidence level, the wider the confidence interval (the most commonly used confidence levels being at 90% or 95%). To foster clarity and comparability it is necessary that the reporting parties, when reporting a CI, clearly specify at least: (1) which error components have been included in the CI; (2) the confidence level and (3) the method used for calculating the confidence interval.

**Figure 3:**
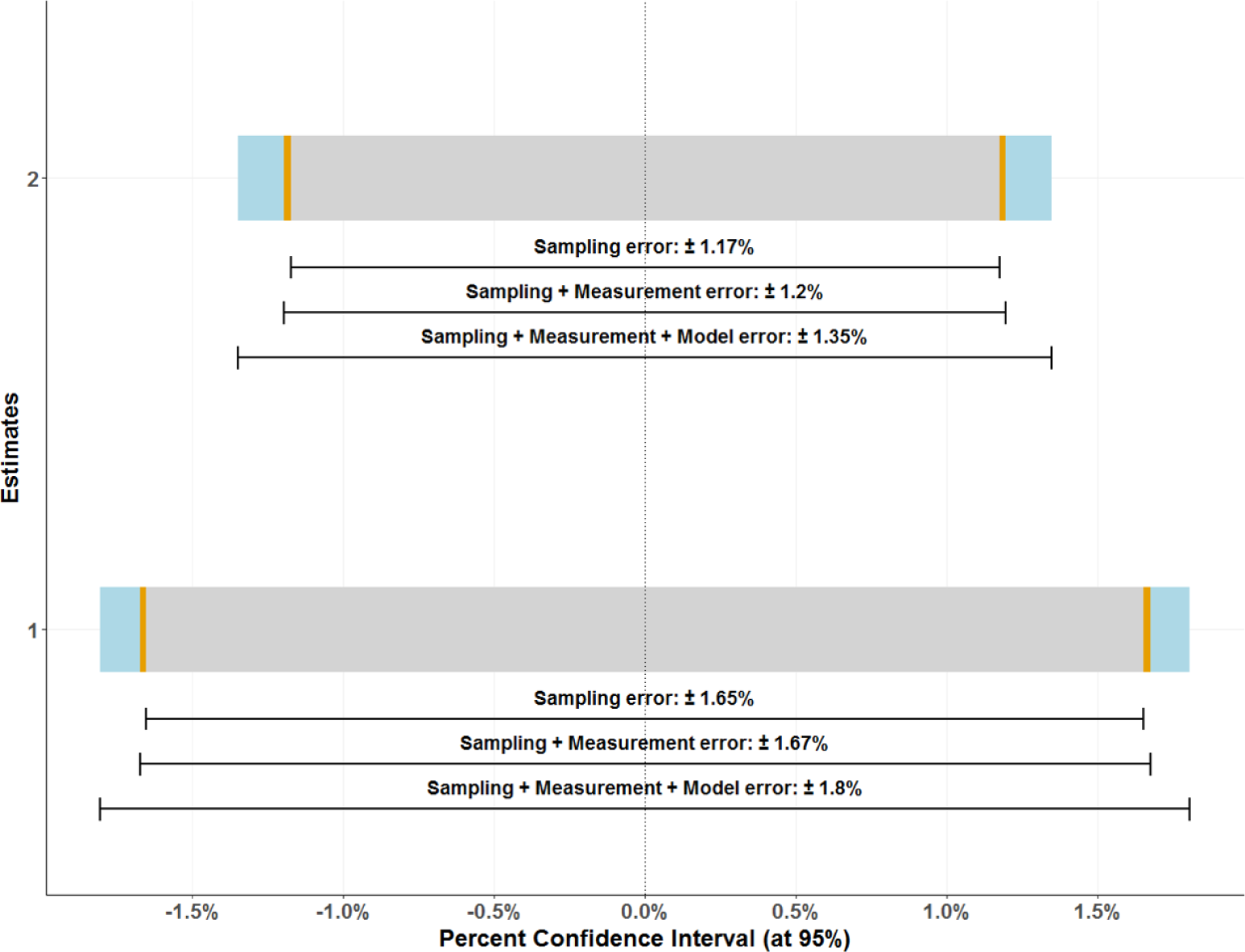
Compounding errors. Graphic representation of confidence intervals (at 95%) of two estimates of average tree volume density in Minnesota (Minnesota Survey Unit 1, USA). Both confidence intervals include three error sources: sampling error (in gray), measurement error (in yellow) and model error (in blue). In both cases the sampling error is the one that contributes the most to the total error. The confidence intervals are derived from data presented in McRoberts et al. (2016, Table 1)

## 3. Conclusions

Estimations of GHG emissions and removals for the LULUCF sector are often obtained using a combination of remotely sensed and ground-based observations. In both cases this information is usually collected through survey sampling techniques. In order to be compliant with the IPCC good practice guidance for greenhouse gas inventories the estimation is required to be transparent, unbiased and as precise as possible. However, assessing the precision and accuracy of large-area surveys is not a easy task, requiring a considerable amount of information on the planning and implementation of the survey, on the data analysis and on the elaboration of the estimates.

In the context of LULUCF reporting, better (i.e., higher quality) uncertainty assessments including more error sources estimates will necessarily bring along wider compounded uncertainties, while systematic errors will be hardly avoided. This apparent absurdity can be rooted in the *uncertainty paradox*, wherein uncertainty aversion determines choices in individuals/institutions (Roeser, 2014). In the current context it would imply that Parties may prefer to report less accurate estimates as long as they present narrower errors. In fact, under UNFCCC (2009) guidance, the paradox is unsolved, since countries are advised to reduce uncertainties through increasing transparency. Technical assessments of Forest Reference Level submissions further advice countries to improve coverage on additional sources of errors (Sandker et al., 2018), which may factually balance the paradox on the side of transparency.

In order to improve the transparency of the reporting, we propose a list of the survey features that should be reported to enable judgment of the quality of the survey results. In this manuscript, the questions proposed for the interrogation of estimates for quality and the sources of error described apply both to ground-based surveys and remote sensing-assisted estimation of activity data. The impact of many sources of errors is often not measurable, difficult to quantify or in the worst case, unknown. Further research is required especially on non sampling errors and techniques to propagate errors in general. It is important for a Party aiming to demonstrate the transparency of its results to explain whether and how they have tackled the main issues that may have an influence on the quality of the estimates.

## Acknowledgements

The authors want to thank specially Becky Tavani and Julian Fox from the Forestry Department in FAO, and the entire UN-REDD program for their comments and continuous support in the preparation of this document.

## Supplementary Materials

### Summary of the key questions

#### General information about the survey

1. Sampled population
  a. What is the population from which the sample is chosen?
  b. If the sampled population is defined as a geographic area, are you able to provide a map of it?
  c. Is the sampled population defined as a finite set of discrete spatial units? If so, which ones? are they uniform? what is their area?
2. Target population
  a. What is the population for which we want to estimate emissions/removals?
  b. For which time period do we need the emissions/removals?
  c. If the target population is defined as a geographic area, are you able to provide a map of it?
  d. If the target population is not defined as a geographic area, are you able to provide a list of the elements that compose it?
3. Sampling selection
  a. Is the survey based on a probability sample?
  b. What sampling design has been used?
  c. What was the planned size of the sample?
  d. What is the size and shape of the sampling units?
  e. Is the sampling unit composed of a cluster of subplots?
  f. Is the sampling unit composed of one or more nested smaller sub-plots?
  g. Was the sample selected following stratified sampling?
  h. If so, which are the strata? how were they constructed? What is their size?
  i. If so, when the strata are defined as geographical areas, are you able to provide a map of them?
  j. Repetition: Is the survey isolated or is it part of a series of repeated surveys? If so, what is the proportion of samples that were repeated?
4. Data collection and processing
  a. Which attributes have been observed in the sampling units?
  b. What was the measurement protocol used to measure the variables of interest?
  c. Has a written field manual or evaluation protocol been produced? Can the party provide it?
  d. Can the party provide a clear and unambiguous definition for each class of categorical variables in the survey, including a land cover classification system (if any)?
  e. Has the classification system (if any) been modified during the implementation of the survey? If so, how does that impact the final estimates? Did the party put in place any system to account for this change?
  f. Has the field measurement protocol been changed during the implementation of the survey? If so, how does that impact the final estimates? Did the party put in place any system to account for this change?
  g. How were the data stored and processed?

#### Information about error sources

5. Sampling error
  a. Which estimator has been used to estimate the emission/removal? Can the party provide the mathematical formula used?
  b. Has the variance of the estimator been estimated? If so, which estimator was used to compute it? Can the party provide the mathematical formula used?
  c. Is the estimator for the population parameter and its variance estimator unbiased (or approximately unbiased) under the sample design adopted in the survey?
6. Model error
  a. Was the variable of interest observed or estimated using a model?
  b. If so, what model was used?
  c. What auxiliary variables have been used to estimate the variable of interest? How have they been collected?
  d. Were the auxiliary variables available for all population elements?
  e. Was the model developed independently of the survey (i.e. selected from models already published in the literature)?
  f. How was the model selected and how can the party ensure that the population for which the model was developed is similar to the target population?
  g. If the data used to develop the model were collected within the survey, can the party provide a description of the survey design and of the model fitting method?
  h. Was the error in model parameters estimated? how?
7. Under-coverage
  a. Is the area over which we want to report the emission/removals entirely included in the area that has been sampled?
  b. If not, how did the party ensure the estimates are representative of whole target population?
8. Over-coverage (domain estimation)
  a. Is the area over which we want to report the emission/removals (the target population) smaller than the area that has been sampled and included in it?
  b. If so, has the party used any domain estimation techniques? Which ones?
9. Non-response
  a. Did the party put in place any measure for the prevention or avoidance of non-response before the data collection? If yes, which ones?
  b. How many of the selected sampling units have proven to be not measurable/not accessible?
  c. Which are the main causes for the non-response? Are there any reason to believe that non-responding elements may be systematically different from the responding ones?
  d. Did the party adjust the estimate to overcome the fact that not all the sampling units have been measured/accessed? If so, how?
10. Time coverage issues
  a. When was the data collection carried out? What is the time period in which the variables of interest were observed?
  b. Is the period over which we want to report emissions/removals included in the data collection period?
  c. If not, how did the party ensure that estimates are representative of the reporting time period?
11. Measurement error
  a. Which measuring instruments were used to measure the variables of interest? Is there any information available about the nominal precision of the measuring instruments used?
  b. Which instruments were used to record the geographical location of the observations? Is there any information available about the nominal precision of the instruments used?
  c. Can the party provide the precision with which each measurement is taken (for example, diameter at breast height to be recorded in cm to the nearest 0.1 cm)?
  d. Did the party put in place any measure for the reduction of the measurement error (including position error)? If yes, which ones?
  e. How many observers/field teams have been employed in the survey?
  f. which satellite products were used to estimate activity data? what are the product specifications?
12. Processing errors
  a. Did the party put in place any measure(s) for the prevention of data entry error in the field and in the office? If yes, which?
  b. Which methods were used for the identification of invalid or aberrant values after the data were entered?
  c. Which methods were used for correcting missing data and outliers?

#### Information about the total error

13. Error propagation
  a. Did the Party propagate errors to establish the total error?
  b. Were all the errors symmetric or normally distributed? If not, how were those errors reported?
  c. Which assumptions were taken in regard to the existence of correlations among the different errors?
  d. If errors were propagated, which method was used and how was it justified?
  e. If the total error followed an asymmetric distribution, how was it reported?

## Footnotes

1 Under a simple random sampling design the variance of the estimator of the sample mean is given by the variance of the population divided by number of elements in the sample.

2 Non-probabilistic sample selection is sometimes carried out in the context of REDD+ and the LULUCF sector. This can happen, for example, whenever the sample is selected based on expert choice (it can be the case of training point selection for supervised land cover classification). As a result, it is not possible to calculate the probability of each population element to be included in the sample (the so-called inclusion probability). Under a design-based inference this results in the fact that sampling variances (and therefore the sampling error) cannot be calculated *unbiasedly*. Conversely, it might not affect the predictions in a model-based framework

